# Complement C1q-dependent engulfment of alpha-synuclein induces ENS-resident macrophage exhaustion and accelerates Parkinson’s-like gut pathology

**DOI:** 10.1101/2023.10.24.563832

**Authors:** PM Mackie, J Koshy, M Bhogade, T Hammoor, W Hachmeister, GM Lloyd, G Paterno, M Bolen, MG Tansey, BI Giasson, H Khoshbouei

## Abstract

Deposition of misfolded α-synuclein (αsyn) in the enteric nervous system (ENS) is found in multiple neurodegenerative diseases. It is hypothesized that ENS synucleinopathy contributes to both the pathogenesis and non-motor morbidity in Parkinson’s Disease (PD), but the cellular and molecular mechanisms that shape enteric histopathology and dysfunction are poorly understood. Here, we demonstrate that ENS-resident macrophages, which play a critical role in maintaining ENS homeostasis, initially respond to enteric neuronal αsyn pathology by upregulating machinery for complement-mediated engulfment. Pharmacologic depletion of ENS-macrophages or genetic deletion of C1q enhanced enteric neuropathology. Conversely, C1q deletion ameliorated gut dysfunction, indicating that complement partially mediates αsyn-induced gut dysfunction. Internalization of αsyn led to increased endo-lysosomal stress that resulted in macrophage exhaustion and temporally correlated with the progression of ENS pathology. These novel findings highlight the importance of enteric neuron-macrophage interactions in removing toxic protein aggregates that putatively shape the earliest stages of PD in the periphery.

## Introduction

Aggregates of α-synuclein (αsyn) are found throughout the enteric nervous system in both Parkinson’s Disease (PD) and Lewy Body Dementia (LBD)^1–5^. It is widely hypothesized that enteric nervous system (ENS) αsyn pathology may precede central nervous system (CNS) pathology in a subset of PD patients, and via prion-like propagation, spreads to vagal fibers and subsequently to the brain^6–10^. Moreover, ENS Lewy pathology is thought to contribute to decreased intestinal motility is found in both PD and LBD leading to severe constipation which can incur high morbidity^11–16^. Thus, enteric synucleinopathy has important implications for pathogenesis and pathophysiology of CNS synucleinopathies.

Several reports have recently shown that gut-seeded αsyn pathology can spread to the brain in animal models^17–20^. These studies have focused on the kinetics, anatomical routes, and modulators of the progression from gut-to-brain. However, relatively little is known about the factors that initially shape the local histopathology and associated functional deficits of enteric synucleinopathy.

The ENS contains specialized macrophages that play important roles in homeostasis, development, and infectious states^21–26^. Given the accepted role for CNS macrophages in the progression of PD pathology^27–29^, it has been hypothesized that these ENS macrophages may play a role in the spread of enteric αsyn aggregates to the CNS^30^. CNS-resident macrophages have been shown to engage in both beneficial and detrimental functions during neurodegenerative diseases and represent promising targets for new therapies^31–33^. Prominent examples include activating the adaptive immune response, which can lead to neuronal death^34–36^, and microglia-mediated complement-dependent synaptic pruning^37–40^, which mechanistically links protein aggregates to neuronal dysfunction. Given the range of states macrophages can adopt, and the disease-modifying potential of macrophage phenotype, it is important to elucidate the precise response of ENS macrophages to synucleinopathy and how this response contributes to development and/or progression of the disease process.

Here we show induction of enteric synucleinopathy disrupts enteric neuron-macrophage interactions and elicits a macrophage response characterized by upregulated synaptic pruning machinery. Macrophages utilize complement to clear αsyn from nearby neurons and constrain the spread of αsyn pathology. However, this mechanism facilitates intestinal dysmotility and eventually leads to macrophage exhaustion as synucleinopathy spreads. Overall, our results reveal a novel mechanism by which ENS-resident macrophages sculpt the histopathological and functional manifestations of enteric synucleinopathy and shed light on potential mechanisms underlying the spreading of peripheral pathology from the gut to the brain in PD.

## Results

### The early ENS macrophage response limits the spread of enteric synucleinopathy

To assess the potential role for the gut-resident immune system in modulating local or distal synucleinopathy, we employed a recently described gut-first model of α-synuclein (α-syn) pathology^17^ by injecting preformed fibrils of mouse recombinant α-syn directly into the stomach and duodenal wall of wild type mice (Fig 1A). We first assessed the time course for local seeding of α-syn pathology within the enteric nervous system using whole mount immunofluorescent microscopy at 4, 14, and 30 days post injection (dpi), and found that enteric synucleinopathy was first observed within the myenteric plexus of pre-formed fibril (PFF)-injected mice at 14dpi, and further increased through 30dpi and 60dpi (Fig 1B,C). The establishment and initial progression of enteric synucleinopathy was observed without changes to total neuronal area (Fig S1A,B). Interestingly, we noticed that macrophages near diseased ganglia directly contacted pSer129+ neurons and contained internalized pSer129+ punctae (Fig 1D). These punctae and the contact between macrophages and neurons was increased in our PFF mice without changes in total macrophage area (Fig 1E-G, and Fig S1C,D). Thus, enteric synucleinopathy alters local macrophage neuroimmune communications.

**Figure 1:**
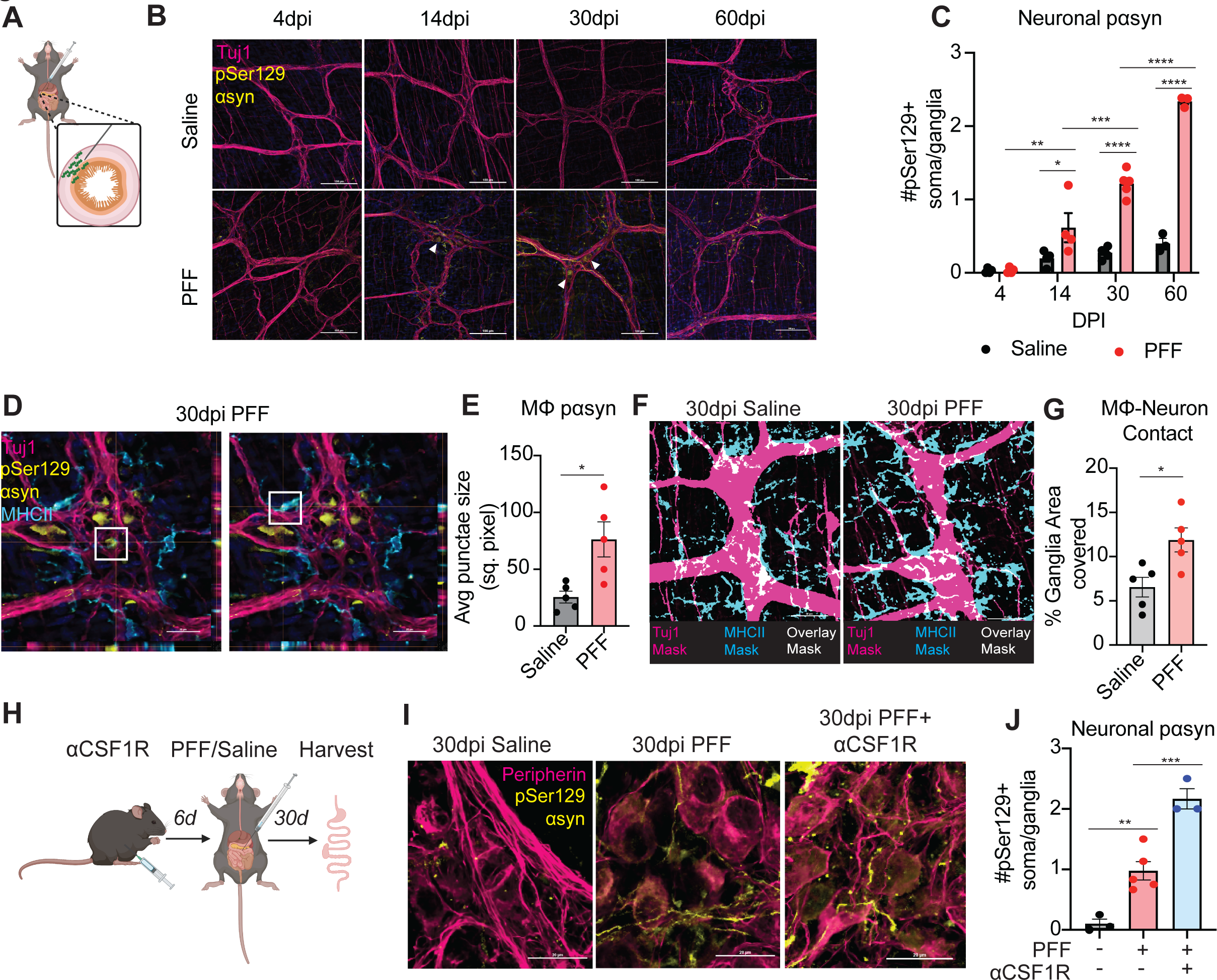
Myenteric macrophages modulate enteric synucleinopathy. (A) Cartoon depicting gut wall injections of PFFs. (B) Representative confocal images of the myenteric plexus (Tuj1, pseudo colored magenta) and pSer129 α-synuclein (pseudo colored yellow) at 4, 14, and 30 days post injection (dpi) with either saline or PFFs. (C) Counts of the average number of pSer129+ neuronal soma/ganglia within the myenteric plexus shows PFF but not saline injection induces neuronal pSer129 pathology starting at 14dpi, with a time-dependent increase to 30dpi. Two-way ANOVA with Tukey’s multiple comparison test: interaction p<0.0001, injection p<0.0001, time p<0.0001; n=4-5 animals/group/time point. (D) Representative confocal image of myenteric plexus (Tuj1, pseudo colored magenta), myenteric macrophages (MHCII, pseudo colored cyan), and pSer129 α-synuclein (pseudo colored yellow) from 30dpi PFF mice depicting a macrophage contacting a pSer129+ soma (left) and a macrophage with internalized pSer129+ punctae (right). (E) Quantification of the average pSer129+ punctae size in myenteric macrophages at 30dpi showing a statistically significant increase in PFF-injected mice. Unpaired t-test, p<0.05; n=5 animals/group. (F) Thresholded and binarized confocal images depicting the contact sites (white) between the myenteric plexus (magenta) and myenteric macrophages (cyan) at 30dpi. (G) Quantification of the average macrophage-neuron contact site size. Unpaired t-test, n=5 animals/group. (H) Cartoon of design for macrophage depletion experiments. (I) Representative confocal images of myenteric ganglia (Peripherin, pseudo colored magenta) with pSer129 α-synuclein pathology (pseudo colored yellow) at 30dpi. (J) Counts of average number of pSer129+ neuronal soma per myenteric ganglia replicating that PFF injection induces neuronal pSer129+ α-synuclein pathology at 30dpi and additionally showing macrophage depletion with PFF injection further increases neuronal pSer129 α-synuclein burden. One-way ANOVA (p<0.0001) with Sidak’s test for multiple comparisons; n=3-5 animals/group.

We next sought to characterize the gut-to-brain spread of α-syn pathology, which has been reported to varying degrees in this model^17–19^. At 30dpi, we did not observe overt pSer129 pathology within the Dorsal Motor Nucleus of the Vagus (DMV) or midbrain dopaminergic nuclei (Fig S1E). We did, however, find an increase in microglial morphologic complexity within the DMV (Fig S1F) as an early indicator of microglial reactivity. Taken together, these data indicate that synucleinopathy confined to the enteric nervous system perturbs myeloid-neuron interactions both locally within the ENS, and distally within the CNS, the latter occurring without bona fide αsyn pathology.

We next inquired whether or not macrophage-neuron interactions have any effect on the development or progression of enteric synucleinopathy. To address this, we depleted peripheral macrophages using injections of a monoclonal antibody against CSF1R^24^, which is required for macrophage survival, without depleting central microglia (Fig 1H, Fig SG, H). Interestingly, loss of ENS macrophages further enhanced the ENS phospho-αsyn pathology induced by PFF injection at 30dpi (Figure 1I, J). Collectively, these data show that, at early stages, myenteric macrophages constrain the development and progression of gut-first synucleinopathy.

### Complement-expressing, antigen-presenting myenteric macrophages are associated with gut synucleinopathy

To determine how macrophages were protective against enteric synucleinopathy, we first assessed various macrophage morphologic parameters, none of which were significantly different (Fig S1I-L). To provide a more sensitive analysis of the macrophage state during enteric synucleinopathy, we performed single-cell RNA sequencing on duodenal immune cells at 30dpi which yielded 27,648 high-quality cells including macrophages, T- and B-lymphocytes, and several non-immune contaminating populations (Fig S2A-C, Table S1). To conduct a more thorough analysis on the macrophage transcriptional response to synucleinopathy, we sub-selected cells enriched for macrophage/myeloid markers *Cx3cr1, Aif1, and Lyz2* and re-clustered them at a higher resolution which yielded 9 transcriptionally distinct macrophage clusters (Fig S2D, Fig 2A,B, Table S1). Analysis for DEGs across all macrophage subsets revealed upregulation of iron-related genes like *Ftl1*, chemotactic genes *Ccl5*, and lysosomal gene *Cd63* in macrophages from PFF-injected mice. Conversely, inflammasome-related genes such as *Nlrp1b* and *Il1r2* were downregulated (Fig 2C, Table S2). Thus, enteric synucleinopathy induces subtle transcriptional changes to the gut macrophage compartment.

**Figure 2:**
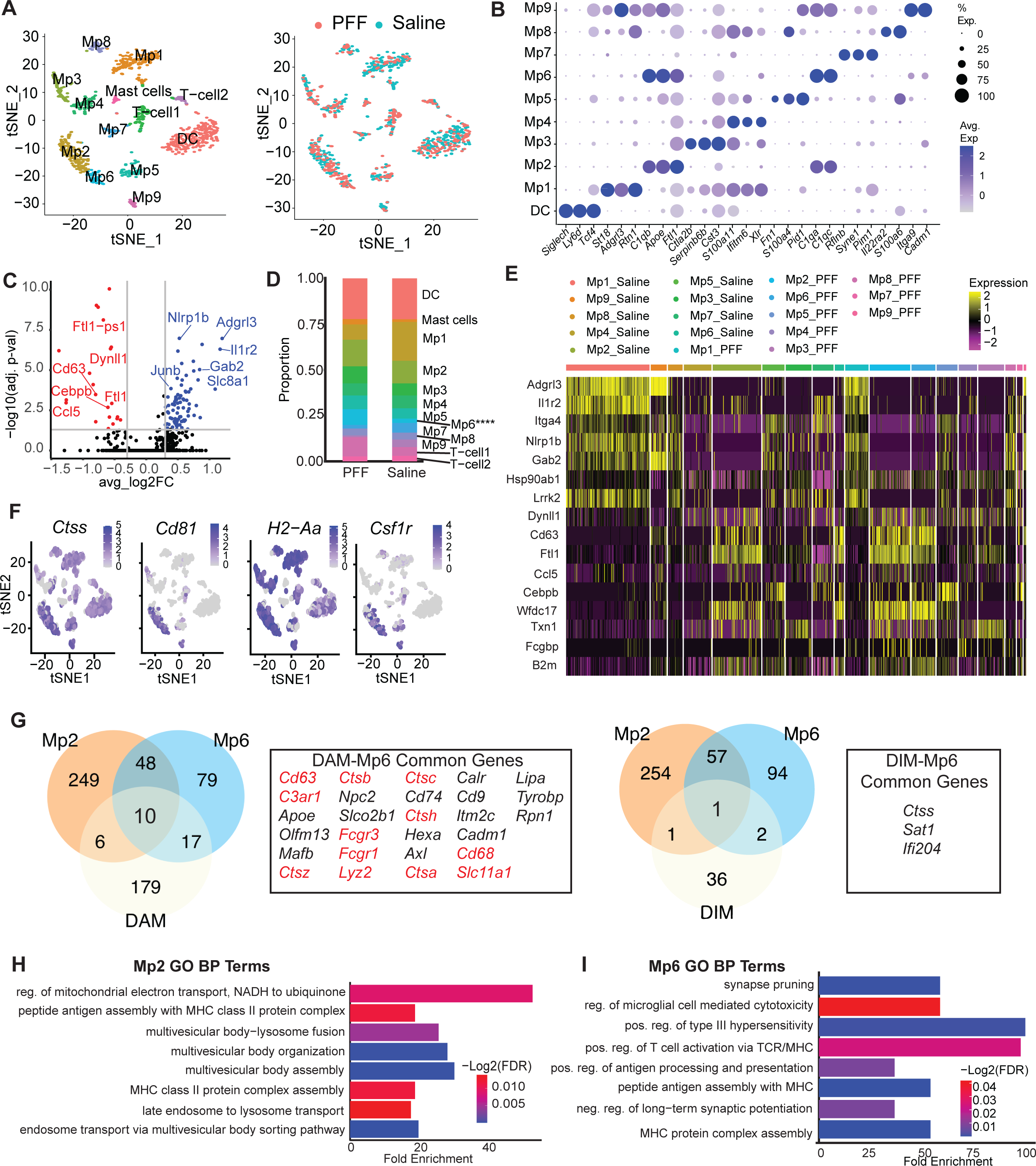
Emergence of a transcriptionally-defined macrophage subset associated with enteric synucleinopathy. (A) t-stochastic neighbor embedding (t-sne) plots of subsetted gut macrophages from saline- and PFF-injected mice at 30dpi showing 9 transcriptionally distinct clusters of macrophages. (B) Dot plot of dendritic cell and macrophage clusters showing the top 2-3 marker genes identifying each cluster. (C) Volcano plot of differentially expressed gene (DEG) analysis of all macrophages between saline control vs. PFF indicates 83 significantly upregulated genes including *Ftl1, Cd63, and Ccl5*, and 21 significantly downregulated genes including *Nlrp1b, Il1r2, and Gab2.* (D) Composition of the PFF cells and Saline cells by cluster indicating that Mp6 was significantly enriched in the PFF group relative to the saline group. Fisher’s exact test, p<0.0001. (E) Heatmap showing expression of the select DEGs identified in (C) across the macrophage clusters. Upregulated DEGs were enriched in Mp2-PFF and Mp6-PFF and, to a lesser extent, Mp2-Saline, and Mp6 Saline. (F) Feature plots depicting the expression of genes previously associated with myenteric macrophages such as *Ctss, Cd81, H2-Aa,* and *Csf1r* across the different macrophage clusters. These genes were highly expressed in Mp6 and in some of the Mp2 macrophages. (G) Venn diagrams comparing the transcriptional signature of the Mp2 and Mp6 macrophages with previously published myeloid cell signatures associated with neurodegeneration such as the disease associated microglia (DAM, left) and the disease inflammatory macrophages (DIM, right). While Mp2 and Mp6 signatures were more similar to each other (58 shared markers) than previously published datasets, but Mp6 did share a larger set of markers with the DAM signature relative to the DIM signature nearly half of which (12/28) were associated with the endo lysosomal system (red text). (H, I) Gene Ontology Biological Process (GO BP) analysis on the significantly expressed marker genes of Mp2 (H) and Mp6 (I) indicating that Mp2 was mainly characterized by enrichment of terms relating to the multi-vesicular body system including processing by the lysosome while Mp6 was mainly characterized by enrichment for antigen-presentation terms and terms relating to synapse elimination including pruning and negative regulation of long-term potentiation. These data were compiled from n=4-6 mice per group.

We then sought to determine if any of the macrophage clusters were specifically associated with synucleinopathy. We analyzed the composition of our saline and PFF groups by cluster and found that Mp6 was significantly enriched in our PFF group (Fig 2D). When we examined the expression of the identified DEGs, we noticed that upregulated DEGs were specifically expressed in clusters Mp2 and Mp6 (Fig 2E). Hence, we noted Mp2 and Mp6 as clusters of interest and labelled Mp6 as our synucleinopathy-associated macrophages (SAMs). Furthermore, the transcriptional signature of Mp2 and Mp6 were highly similar. Not only did they both express the upregulated DEGs, but they also both expressed high levels of C1q components, *Apoe*, and antigen presentation genes such as *H2-Aa* and *Cd81* (Fig 2B,F). The latter are consistent with previous observations of conserved MHCII^hi^ macrophage interstitial populations across tissues^41^ and the high MHCII expression of myenteric macrophages^24^. Mp2 and Mp6 also highly expressed *Csf1r*, which is crucial for the survival and phenotype of ENS-resident macrophages^24^, and the canonical microglial gene *Ctss*, which is also highly expressed in nerve-associated macrophages^23^ (Fig 2F). Therefore, it is highly likely that our Mp2 and Mp6 clusters represent ENS macrophages.

We then asked how SAMs related to previously published myeloid cell phenotypes in the context of neurodegenerative disease. So, we compared our Mp2 and Mp6 populations to the well-established DAM signature^42^ and the recently described Disease Inflammatory Macrophages (DIM) signature^43^. 27 genes were shared between our SAM (Mp6) and the published DAM signature, many of which were lysosomal proteases such as cathepsins that have been implicated in α-syn clearance^44, 45^, compared to only 3 shared between the SAM and DIM signatures (Fig 2G). These data highlight the similar transcriptional responses of nervous-system resident (central and enteric) myeloid cells across different proteinopathies and organs and underscore the importance of lysosomal function in disease process.

Finally, we functionally annotated Mp2 and Mp6 using Gene Ontology. Our Mp2 cluster was highly enriched in terms related to the multivesicular body-lysosome pathway, which has been implicated in α-syn aggregation^46^, whereas Mp6 was highly enriched in antigen presentation terms, synaptic pruning, and negative regulation of long-term potentiation (Fig 2H,I). Enrichment of antigen-presentation terms were corroborated by our T-cell sequencing dataset that showed upregulation of genes relating to T-cell activation and exhaustion in PFF-injected mice (Fig S2E-G). The synaptic pruning term was corroborated by high expression of C1q genes, *C3*, and early complement component receptors in Mp6 macrophages (Fig 2B, Table S1). Overall, these data revealed an α-synucleinopathy-associated macrophage cluster that was defined by high expression of complement-, antigen-presentation-, and lysosomal genes all of which have been previously associated with α-syn pathology.

### Complement-dependent engulfment of α-syn by SAMs couples enteric pathology and dysfunction

We were particularly interested in the synaptic pruning term highlighted in our GO analysis of Mp6 macrophages. We first wanted to validate the C1q expression indicated by our scRNA-seq on a protein and spatial level, as C1q genes were most highly expressed by Mp6 and, to a lesser extent, Mp2. Using whole-mount immunofluorescence, we confirmed that ENS macrophages express C1q protein and that expression is increased in PFF-injected mice (Fig 3A,B). These data are consistent with a recent report that macrophages are the main sources of C1q within the intestine^47^. Interestingly, we observed C1q+/pSer129 α-syn+ punctae within ENS macrophages, and these were also increased in PFF-injected mice (Fig 3C) indicative of complement-mediated internalization of α-syn by ENS macrophages. These findings were associated with increased deposition of C1q on myenteric neurons (Fig 3D,E), which, taken together, suggest that myenteric macrophages utilize complement to engulf αsyn from enteric neurons.

**Figure 3:**
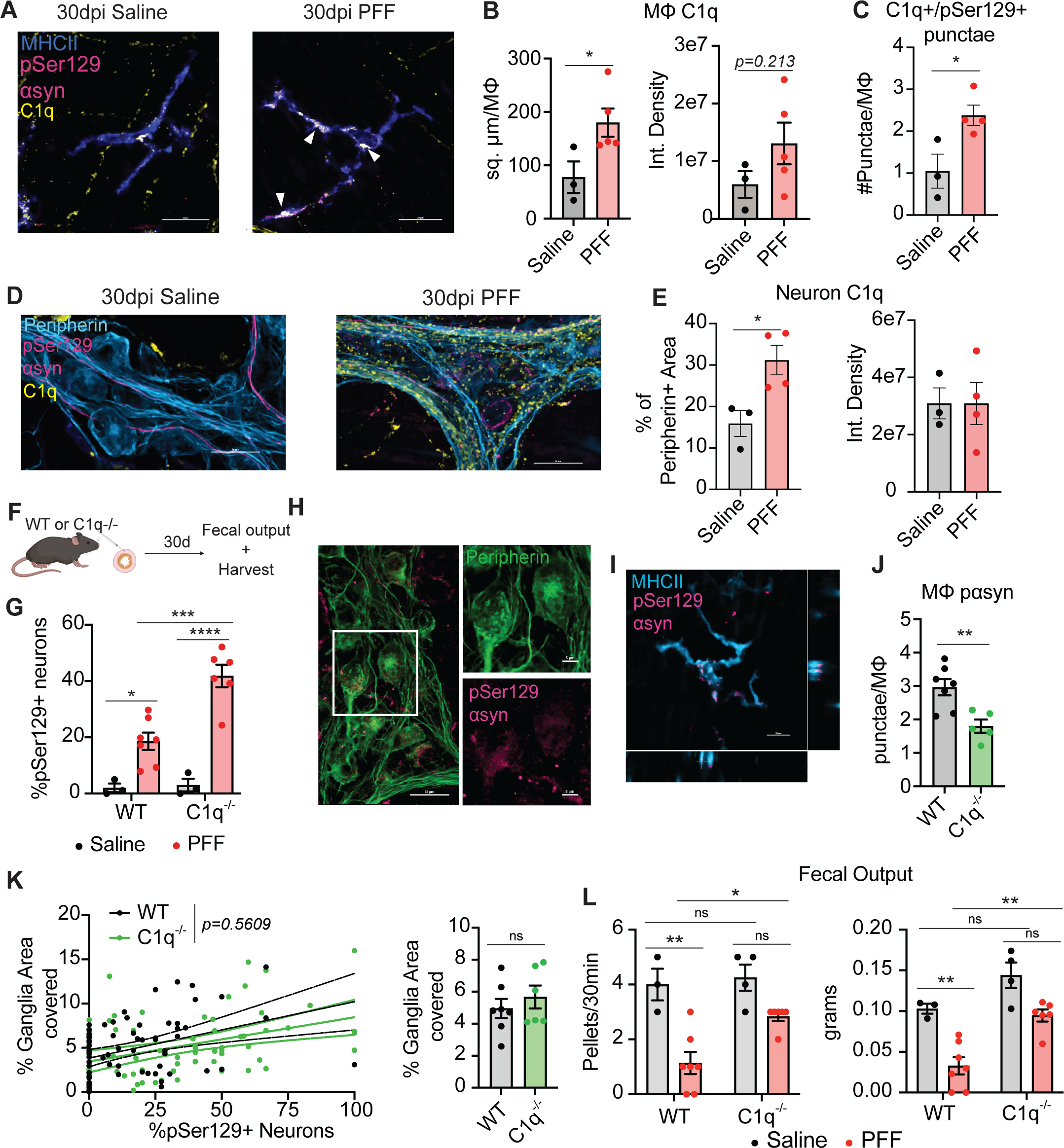
ENS-resident macrophages engulf α-syn from neurons via a C1q-dependent mechanism. (A) Representative confocal images of myenteric macrophages (MHCII, blue) at 30dpi saline or PFFs showing expression of C1q (magenta) and co-localized punctae of C1q with pSer129 α-syn (magenta, white arrowheads) in PFF mice. (B) Quantification of C1q+ area and intensity within myenteric macrophages showing upregulation in PFF-injected mice Mann-Whitney test, p<0.05, n=3-5 animals/group). (C) Quantification of the number of C1q+/pSer129 α-syn+ punctae within myenteric macrophages showing an increase in PFF mice (unpaired t-test, p<0.01, n=3-5 animals/group). (D) Representative confocal images of myenteric ganglia (peripherin, cyan) at 30dpi with either saline or PFF showing pSer129 α-syn+ soma (yellow), and deposition of C1q+ punctae (magenta) on neuronal soma. (E) Quantification of the C1q+ area (left) and intensity (right) on myenteric neurons in saline and PFF mice showing an increase with PFF injection (unpaired t-test, p<0.05, n=3-4 animals/group). (F) Cartoon depicting experimental design for C1qa^-/-^ experiments. (G) Quantification of the number of pSer129 α-syn+ neuronal soma expressed as a percentage of total peripherin+ soma within the myenteric plexus of saline- and PFF-injected mice at 30dpi in both C1q^-/-^ and WT controls showing a PFF-induced increase in pathology that was further enhanced in the C1q^-/-^ mice (two-way ANOVA p_condition_<0.0001, p_genotype_<0.01, p_interaction_<0.05. Tukey’s test for multiple comparisons WT-saline vs WT-PFF p<0.05, WT-saline vs C1q^-/-^-saline p=0.9987, C1q^-/-^-saline vs C1q^-/-^-PFF p<0.0001, WT-PFF vs C1q^-/-^-PFF p<0.001). (H) Representative confocal images of myenteric ganglia (peripherin) showing qualitatively increased pSer129 α-syn+ (color) neurons in C1q^-/-^-PFF animals. (I) Representative confocal images of myenteric macrophages (MHCII, color) with internalized pSer129 α-syn+ punctae (color, white arrowhead) in PFF injected C1q^-/-^ mice and WT controls 30dpi. (J) Quantifcation of the number of internalized pSer129 α-syn+ within MHCII+ myenteric macrophages showing decreased punctae in C1q^-/-^ macrophages (unpaired t-test, p<0.01). (K) Linear regression analysis of 139 myenteric ganglia comparing the percentage of pSer129+ neurons against the neuron-macrophage contact showing a statistically significant positive correlation that did not differ between genotypes (p=0.5609, left). Average contact size between macrophages and myenteric ganglia in PFF-injected WT and C1q^-/-^ mice revealing no significant difference between groups (unpaired t-test, right). (L) Fecal output measured by the number of pellets or the total weight of pellets after a 30-minute period showing a decrease in the PFF group that was partially rescued by loss of C1q. Pellet count (left): two-way ANOVA p_condition_<0.001, p_genotype_<0.05 with Tukey’s test for multiple comparisons WT-saline vs WT-PFF p<0.01, WT-saline vs C1q^-/-^-saline p=0.9819, C1q^-/-^-saline vs C1q^-/-^-PFF p=0.100, WT-PFF vs C1q^-/-^-PFF p<0.05. Pellet weight (right): two-way ANOVA p_condition_<0.0001, p_genotype_<0.001 with Sidak’s test for multiple comparisons WT-saline vs WT-PFF p<0.01, WT-saline vs C1q^-/-^-saline p=0.2319, C1q^-/-^-saline vs C1q^-/-^-PFF p<0.05, WT-PFF vs C1q^-/-^-PFF p<0.01.

To validate that complement dependent engulfment α-syn was a protective mechanism against enteric synucleinopathy, we repeated our gut PFF injections in C1qa^-/-^ mice and WT controls (Fig 3F). Loss of C1q resulted in a further increase in PFF-induced enteric neuronal pSer129 α-syn+ staining compared to WT controls (Fig 3G, H). The exacerbated neuronal pathology was accompanied with a decrease in the number of internalized pSer129 α-syn+ punctae within macrophages (Fig 3I, J), supporting interpretation that C1qa^-/-^ mice were less efficient at clearing α-syn from neurons, leading to spread of pSer129+ pathology. Consistent with our previous analysis (Fig 1F,G), there was a positive correlation between the neuronal pSer129 α-syn+ pathology load and the amount of neuron-macrophage contact that was unaffected by loss of C1q (Fig 3K), suggesting that chemotactic signaling remained intact.

Complement-dependent engulfment has frequently been implicated in excessive synaptic elimination leading to functional deficits^37, 38, 40^. In the CNS, α-syn is known to be enriched at presynapses^48^, but the subcellular localization of α-syn within enteric neurons is unknown. As predicted, we found that enteric α-syn highly co-localized with pre-synaptic but not post-synaptic markers (Fig S3). We therefore examined the role of C1q in enteric neuronal function. It has previously been shown that fecal output is a sensitive assay for gross intestinal dysmotility in this animal model of synucleinopathy^18^. Indeed, we found decreased fecal output in our PFF-injected mice relative to saline controls that was partially rescued by loss of C1q (Fig 3L). Thus, loss of C1q decoupled enteric neuron histopathology and functional deficits, indicating that pSer129 pathology induced enteric dysfunction in part via a complement C1q-dependent mechanism.

### ENS macrophages adopt an immune-exhaustion phenotype as gut synucleinopathy progresses

Thus far, our data indicate that myenteric macrophages upregulate synaptic pruning machinery to engulf αsyn from neurons thereby limiting the spread of synucleinopathy in the gut at early time points. Given that our data demonstrated further progression of pSer129 α-syn+ pathology from 30 to 60dpi (Fig 1B, C), we asked if this macrophage response was sustainable or if it eventually failed. We repeated our PFF injections and waited up to 60dpi then re-examined the enteric nervous system for αsyn pathology and C1q. While macrophages at 60dpi maintained a comparable level of C1q expression relative to 30dpi, they contained fewer internalized C1q+/pSer129+ punctae (Fig 4A, B). There was also decreased C1q deposition on enteric neurons at 60dpi compared to 30dpi (Fig 4C, D). Collectively, these data suggest that ENS macrophages lose their C1q-dependent engulfment capacity as enteric synucleinopathy progresses.

**Figure 4:**
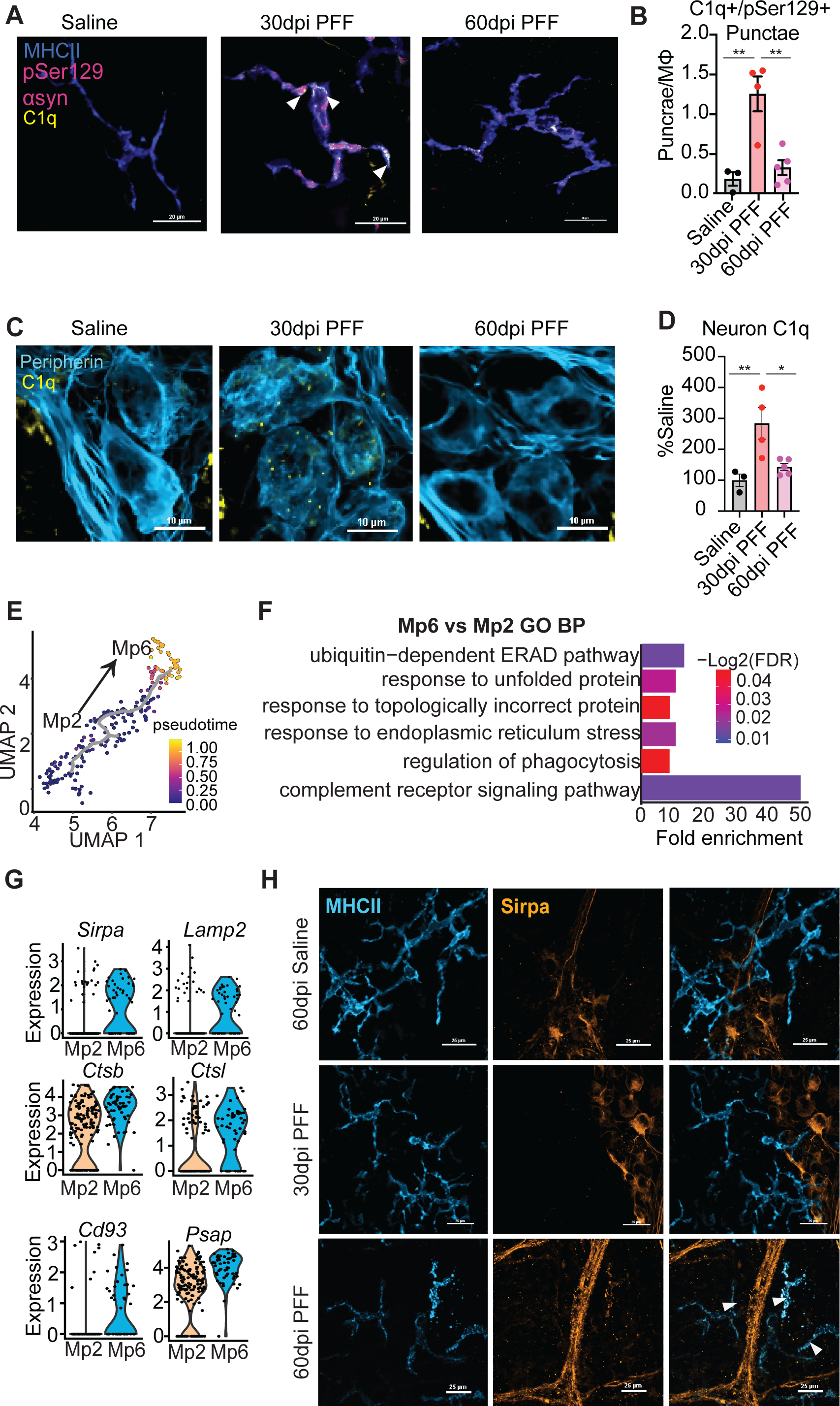
ENS synucleinopathy gradually exhausts macrophage clearance capacity. (A) Representative confocal images of ENS-resident macrophages (MHCII, blue), pSer129 α-syn (magenta), and C1q (yellow). Arrow heads indicate co-localized pSer129+/C1q+ punctae within the macrophage. (B) Quantification of the number of pSer129+/C1q+ punctae per macrophage showing an initial increase at 30dpi that then decreases by 60dpi. Ordinary one-way ANOVA (p<0.01) with Sidak’s multiple comparisons test (**p<0.01). (C) Representative confocal images of myenteric neuronal soma (Peripherin, cyan) and C1q (yellow) showing complement deposition at 30dpi PFF. (D) Quantification of C1q deposition (yellow) on peripherin+ area (cyan) shown as % of saline control. C1q deposition increased at 30dpi PFF and then decreased to near-control levels at 60dpi. Ordinary one-way ANOVA (p<0.01) with Sidak’s test for multiple comparisons (*p<0.05, **p<0.01). (E) Trajectory analysis of Mp2 and Mp6 clusters using Monocle v3.0 shows differentiation from Mp2 to Mp6 through pseudo-time. (F) Gene Ontology analysis of significant DEGs between Mp2 and Mp6 showing upregulation of terms related to endo-lysosomal stress, ER stress, and the unfolded protein response in Mp6 macrophages. (G) Violin plots of select DEGs significantly upregulated in Mp6 vs Mp2 including mostly lysosomal genes such as Lamp2 and cathepsins, and Sirpa. (H) Representative confocal images of ENS-resident macrophages (MHCII, cyan) at 30dpi and 60dpi showing expression of Sirpa (orange) at 60dpi.

We then investigated the mechanism responsible for the loss of macrophage clearance capacity by examining the transcriptional relationship between Mp6 and Mp2. While these clusters had high transcriptional overlap, only Mp6 was associated with synucleinopathy. First, we used Monocle to conduct a trajectory analysis which indicated that Mp2-like macrophages can differentiate into Mp6-like macrophages (Fig 4E). This differentiation was marked by upregulation of intracellular stress related pathways such as the response to topologically incorrect protein, response to ER stress, and the complement receptor signaling pathways (Fig 4F). Additionally, this transition was marked by increased expression of lysosomal genes such as *Lamp2, Psap, Ctsb, and Ctsl* (Fig 4G). Notably, *Ctsb, Ctsl,* and *Psap* have all been implicated in the proteolytic processing of αsyn and are heavily implicated in PD-related pathology^45, 49, 50^. Taken together, these data indicate that myenteric macrophages experience increased endolysosomal stress as they engulf pathologic αsyn. Furthermore, Mp6 macrophages upregulated expression of Sirpa (Fig 4G), a negative regulator of synaptic engulfment^51, 52^, indicative of exhaustion^53, 54^. Additionally, we found Sirpa+ macrophages near myenteric ganglia at 60dpi, but not 30dpi *in situ* (Fig 4H) indicating that macrophage exhaustion temporally correlates with the progression of pSer129 α-syn+ pathology in the ENS. Collectively, our microscopy and sc-RNA sequencing data indicate that enteric synucleinopathy gradually exhausts ENS macrophages by inducing endolysosomal stress, resulting in downregulation of functional clearance capacity thereby facilitating the spread of α-syn pathology in the ENS.

## Discussion

The current understanding of PD has evolved to recognize the importance of ENS involvement both in the pathogenesis of the disease and in disease progression which significantly compromises quality of life for the patient. A major challenge has been elucidating the cell-to-cell interactions and molecular processes that shape the course of enteric synucleinopathy within the context of PD-like pathology. Here, we found that ENS resident macrophages initially engage a protective phenotype characterized by complement-dependent engulfment of α-syn from enteric neurons. Furthermore, our data suggest that while this mechanism limits the histopathological spread of α-syn, it simultaneously contributes to ENS dysfunction. Specifically, the engulfment of toxic α-syn eventually leads to increased endolysosomal stress and ENS-resident macrophage exhaustion which impacts ENS health. Collectively, our data highlight complement-dependent engulfment as a dynamic mechanism by which ENS-resident macrophages modify the complex and multiple aspects of enteric synucleinopathy.

### The complex and multi-dimensional macrophage response to PFFs and synucleinopathy

Macrophages such as microglia can adopt a multitude of states and phenotypes in different neurodegenerative diseases^55^. In PD specifically, several groups have shown that microglia or other CNS-associated macrophages engage in T-cell recruitment^35, 56^ and inflammasome activation^57–59^ to drive neurodegeneration. A previous study found upregulation of multiple pro-inflammatory cytokines in the gut 7 days following PFF injection^18^, suggesting that perhaps ENS macrophages engage a similar pro-inflammatory program to microglia in response to α-syn. In contrast to these findings, we found downregulation of inflammasome-related genes in PFF macrophages. A potential explanation is that the macrophage response during fibril clearance is distinct from that associated with developing neuronal pathology. This agrees with our data indicating macrophages become exhausted over time, suggesting that their response to enteric synucleinopathy is highly dynamic. Characterizing the full trajectory of their response will be critical for the design and timing of any therapeutic intervention.

Instead, our synucleinopathy-associated macrophages were characterized by upregulation of synaptic pruning-related genes and antigen processing/presentation genes. While we focused on the former for this study, it is crucial to recognize the role the adaptive immune system may play in PD progression. Indeed, our sequencing data suggested shifts in the intestinal lymphocyte compartment of PFF-injected mice. Multiple studies indicate that α-syn reactive T-cells are found in the circulation of PD patients^60–62^. Lymphocytes can be educated in the gut before travelling to the CNS^63–65^, and a recent report by the Hong Lab demonstrates that ENS macrophages license T-cells which migrate to the CNS. Thus, it is likely that the C1q-depdendent mechanism of α-syn engulfment we describe is the first step in antigen processing for downstream T-cell activation. More work is required to de-couple these pathways and balance the detrimental and beneficial functions of ENS macrophages.

Endolysosomal stress is thought be a convergence point for multiple pathophysiologic processes across both idiopathic and familial PD^66^ and is thus the subject of intense research. Our data suggest that continued engulfment of toxic α-syn upregulates pathways related to endolysosomal stress leading to exhaustion in ENS macrophages. While it may be tempting to target exhaustion via SIRPa inhibition to promote continued clearance for example, it is highly likely that continued engulfment may induce deleterious responses associated with increased endolysosomal stress, such as inflammasome activation or pyroptosis^67–69^. Further investigation is needed to identify mechanisms to enhance macrophage breakdown of pathologic α-syn species.

### Mechanisms of ENS dysfunction in synucleinopathy

Our results indicate that loss of C1q increases enteric α-syn pathology but paradoxically improves dysmotility. A likely explanation would be that ENS macrophages preferentially engulf α-syn from enteric synapses, thereby limiting the trans-synaptic propagation of misfolded α-syn while simultaneously decreasing neurotransmission efficiency. Indeed, we found that enteric α-syn preferentially localizes to the pre-synaptic space. Moreover, although excessive synaptic pruning due to α-syn has not been shown, it is a well-studied mechanism for cognitive dysfunction in Alzheimer’s-related pathology^37, 38, 70^. Additionally, ENS macrophages engage in synaptic refinement during development^23^. Thus, to our knowledge, our findings are the first to support a role for macrophage-mediated synaptic pruning in response to neuronal α-syn pathology. Notably, increased complement deposition has also been reported in the Substantia Nigra of PD patients^71^. Future studies may investigate whether or not microglia may also engage in synaptic pruning in the context of α-syn pathology and elucidate how macrophages determine which synapses to target for engulfment. Finally, the divergence of pathology and function in our studies is similar to some studies with Aβ^31^. This de-coupling underscores the importance of examining both pathologic and functional outcomes for new interventions.

### Limitations of the model and study design

Here we employed multiple orthogonal approaches including sequencing, pharmacologic depletion, genetic knockouts, and imaging, to describe a novel role for myenteric macrophages in response to synucleinopathy. However, there are still outstanding limitations that future work should address. First, we utilized injections of preformed fibrils to induce synucleinopathy and studied time points focused on early spread of pathology, this model cannot address how synucleinopathy begins. Complementary models using infection, toxins, or inflammation should be employed to determine how the macrophage response to α-syn changes with distinct initial insults. Furthermore, fibrils are a finite nidus and may be taken up by macrophages upon injection. To ensure that the macrophage response characterized here is truly due neuronally-derived α-syn, a complementary model using viral overexpression of α-syn will be informative.

One of our experiments utilized pharmacologic depletion of macrophages via a monoclonal antibody against the mCSF1R receptor. While this technique is standard, it only provides a transient depletion. Indeed, by the 30dpi timepoint, the myenteric niche had been repopulated. Therefore, more sustained and selective modalities of depletion or, alternatively, varying the time of depletion, will be important to corroborate our results. Still, our experiment using C1qa^-/-^ mice provide complementary evidence for our conclusions.

Finally, our study focused on the enteric phase of the gut-to-brain hypothesis of PD. We did not observe any α-syn pathology in the brain at the time points we investigated. Notably, various groups have reported differing time scales and requirements for the gut-brain spread in the PFF model. Therefore, more work is required to standardize this model and rigorously characterize the kinetics of α-syn pathology along the gut brain axis.

In conclusion, we found a novel role for complement-dependent, ENS-resident macrophage engulfment of α-syn in the development and initial spread of enteric synucleinopathy thus introducing a potential therapeutic avenue that may impact that earliest stages of Parkinson’s Disease.

## Acknowledgements

We’d like to thank Dr. Beth Stevens and her lab for providing the anti-C1q antibody. We’d like to thank Dr. Stefan Prokop for his insight and guidance in interpreting neuropathology. We’d also like to thank the Flow Cytometry Core at the University of Florida for their assistance with FACS, and 10x Genomics for their technical support for the single-cell sequencing. Finally, we’d like to thank all members of the Khoshbouei Lab for their thoughtful discussions and contributions throughout the course of this paper. This work was funded from NIH/NINDS grants R01NS071122 (HK), R21NS133384 (HK and BIG), F30NS129283 (PMM), T32-NS082128 (PMM), 1RF1NS28800 (MB and MGT), NCATS TL1TR001428 and UL1TR001427 (PMM), and the Karen Toffler Trust (PMM). We also thank the joint efforts of The Michael J. Fox Foundation for Parkinson’s Research (MJFF) and the Aligning Science Across Parkinson’s (ASAP) initiative. MJFF administers the grant ASAP-020527 on behalf of ASAP and itself. For the purpose of open access, the author has applied a CC-BY public copyright license to the Author Accepted Manuscript (AAM) version arising from this submission.

## Author Contributions

Conceptualization: PMM, MGT, BIG, and HK

Experimental Design: PMM, MGT, BIG, and HK

Data Collection: PMM, JK, MB, TH, WH, GML, GP, and MB

Data Analysis: PMM, JK, MB, WH, GML, GP, and MB

Writing manuscript (original draft): PMM

Revising and editing: PMM, MGT, and HK

