## Supplemental Methods and Figures for "Complement C1q-dependent engulfment of alpha-synuclein induces ENS-resident macrophage exhaustion and accelerates Parkinson’s-like gut pathology"

##### Mice

All mice were used in accordance with national and institutional guidelines and under protocols approved by the University of Florida IACUC. C57B6/J mice were either purchased from Jackson Labs (animals for surgery) or bred in-house (validation experiments). C1qa knockout mice were also purchased from Jackson Labs (strain #031675, RRID IMSR_JAX:031675) at 2-3 months of age. Knockout of C1q was additionally validated in-house by immunofluorescent microscopy.

##### Macrophage Depletion

Macrophage depletion was performed based on the protocol established by Muller et al[1]. Mice were intraperitoneally injected with and a monoclonal antibody against CSF1R (InVivoMab BioXCell, see table for clone) at a dose of 37.5µg/kg. Surgery was performed 6 days after injection.

##### Preformed α-Synuclein Fibrils (PFFs) Generation

Recombinant mouse full length α-synuclein (αSyn) protein was expressed in BL21 (DE3)/RIL Escherichia coli (E. coli; New England BioLabs Inc) using the pRK172 bacterial expression vector containing the murine αSyn cDNA. Bacterial were harvested by centrifugation and re-suspended in high-salt buffer (0.75 M NaCl, 50 mM Tris, pH 7.4, 1 mM EDTA) containing a cocktail of protease inhibitors and heated to 100$^{o}$C for 10 min. Debris were removed by sedimentation at 20,000xg for 30 min. Supernatants were dialyzed into 100 mM NaCl, 10 mM Tris, pH 7.5 and applied onto a Superdex 200 gel filtration column (Amersham Pharmacia Biotech, Inc., Piscataway, NJ) and separated by size exclusion. The fractions were assayed for the presence of the α-Syn proteins by SDS-polyacrylamide gel electrophoresis (PAGE) followed by Coomassie Blue R-250 staining. Proteins were concentrated using Amicon Ultra-15 units (Millipore Corp., Bedford, MA), dialyzed against 10 mM Tris, pH 7.5, applied to a Resource Q column (GE Healthcare) and eluted with a 0-0.5 M NaCl gradient. Protein concentrations were determined using the bicinchoninic acid (BCA) protein assay (Pierce, Rockford, IL) and bovine serum albumin (BSA) as a standard. The purified αSyn monomers were then assembled into PFFs by incubation at 5 mg/mL in sterile PBS at 37 °C with continuous shaking at 1050 rpm for at least 48 hours using an Eppendorf Thermomixer. αSyn fibril formation was monitored with K114 fluorometry as previously described[2]. PFFs were diluted in sterile PBS to a final concentration of 2mg/ml and fragmented into an array of shortened fibrils using a Branson 2800 water bath sonicator at room temperature for 1 hours prior to freeze-down. Once fragmented, PFFs were aliquoted and stored at -80°C until use. At the time of surgery one aliquot was thawed and used then discarded. All surgeries for this study were conducted with PFFs from the same preparation to limit batch effects.

##### Gut Injection Surgery

Mice approximately 3 months old were used for surgery. The gut injections were performed as previously described[3, 4]. Mouse anesthesia induction and maintenance was achieved with isofluorane at 1.5-3%. The abdomen was shaved and sterilized using chlorhexidine/ethanol swabs and a midline incision was made from the xiphoid process to the lower abdomen. Once the peritoneum was opened, the stomach and duodenum were teased out using wetted cotton tip applicators to minimize trauma. Gut injections were performed with a Nanofil syringe fitted with a 36G beveled tip (World Precision Instruments). 4 injections in total were administered to the gut wall: 1 in the stomach antrum, 1 near the pylorus, 1 in the first segment of the duodenum and 1 in the second segment of the duodenum. Each injection was 2.5µl of 2mg/ml PFFs. Fast Green FCF dye (Sigma Aldrich) was added to either the PFFs or sterile saline (control) to visually confirm injection into the gut wall rather than the gut lumen. Following confirmation of injection, the mice were closed and allowed to recover. Mice were housed independently for the first 7-10 days following surgery to allow for the abdominal wound to heal before being recombined with their littermates.

**Duodenal Whole Mount Immunofluorescence**

At designated time points, mice were euthanized via isofluorane overdose followed by transcardial perfusion with cold PBS. The duodenum was extracted and for the remainder of the microdissection was kept submerged cold PBS on ice. The duodenum was flushed intra-luminally with PBS, and the mesentery was gently removed using fine dissection. The duodenum was opened along the mesenteric line and then pinned to a cured Sylgard disc with the mucosa side facing up. The mucosa was then gently dissected off using fine dissection technique under a microscope. Following dissection, the samples were drop-fixed in 4% paraformaldehyde for 20 minutes and then hand-washed in PBS 5 times. Dissected samples were then photobleached overnight and were then ready for immunostaining.

Duodenal samples were first blocked and permeabilized in 0.5% Triton-X, 3% BSA (Sigma), 3% normal goat serum (NGS) for 3 hours at 37°C. Primary antibodies were diluted in 0.5% Triton-X, 2% BSA, 2% NGS. For specific antibodies and dilutions, see Table. Primary labelling incubated overnight with rocking at 4°C. Following primary labelling, samples were washed in PBS 5 times at room temperature for 15 minutes each with rocking. Secondary antibodies (see Table) were diluted in the same solution as primary antibodies and allowed to incubate with samples at room temperature for 2 hours with rocking. Samples were then washed again 4 times for 15 minutes each, counterstained with 4',6-diamino-2-phenylindole (DAPI) at a final dilution of 1:2000 for 20 minutes at room temperature and briefly given one final PBS wash. Samples were then mounted using Fluormount G, sealed, and kept for imaging.

**Mouse Brain Immunostaining**

*FFPE Brains*

Mice were euthanized and transcardially perfused with cold PBS as above and the brains were removed and fixed in 70% ethanol, 150mM NaCl for several days. Brains were sectioned into 8 segments and dehydrated through a series of 70-100% ethanol, followed by xylenes and then infiltrated with paraffin. Paraffinized brains were then embedded in blocks and cut in 5 μm sections mounted on glass slides. Immunostaining was performed using established methods[5]. Briefly, slides were rehydrated by immersing in xylenes, then in graded 100%-70% ethanol steps, followed with heat-induced epitope retrieval (HIER) in a steam bath for 60 minutes in water with 0.05% Tween-20. After antigen retrieval, sections were washed for 15 minutes in running deionized water. Endogenous peroxidase was quenched by incubating sections in 1.5% hydrogen peroxide/0.005% Triton-X-100 diluted in PBS, pH 7.4 for 15-20 minutes. Sections were then rinsed in running deionized H2O for 15 minutes, washed three times for five minutes in 0.1M Tris, pH 7.6, and then blocked in 2% fetal bovine serum (FBS)/0.1M Tris, pH 7.6 solution for five minutes. Slides were incubated with primary antibodies diluted in blocking solution and stored overnight in 4°C.

After overnight incubation, primary antibody was removed from slides with a quick rinse with deionized H2O, then incubated with agitation for five minutes in 0.1M Tris, pH 7.6, three times. Tissue sections were incubated for one hour with anti-mouse biotinylated IgG (Vector Laboratories; Burlingame, CA) in 0.1M Tris, pH 7.6/2% FBS (1:3000) at room temperature. Secondary antibody was rinsed three times with 0.1M Tris, pH 7.6 for five minutes. Sections were then incubated with an avidin-biotin complex (ABC) solution (Vectastain ABC Elite kit; Vector Laboratories, Burlingame, CA) for one hour at room temperature, then rinsed again, three times, with 0.1M Tris, pH 7.6, for five minutes. Sections were developed using chromogen 3,3′-diaminobenzidine (DAB kit; KPL, Gaithersburg, MD) and counterstained using hematoxylin (Sigma Aldrich, St. Louis, MO).

IHC sections were digitally scanned using an Aperio ScanScope CS instrument (40× magnification; Aperio Technologies Inc., Vista, CA, USA), and images of representative areas of pathology were captured using the ImageScope software (4 - 20× magnification; Aperio Technologies Inc.).

*Free-floating immunostaining*

Mice were perfused as described above and the brain was extracted and allowed to post-fix overnight in 4% paraformaldehyde at 4°C. 40 micrometer sections of the Dorsal Vagal Complex (DVC) and midbrain were cut using a vibratome (Leica). To prepare for the cutting procedure, the brain was cut at the optic chiasm and the caudal part, containing the DVC and midbrain, was suspended in 2% agarose and mounted on the vibratome. The brain was cut caudal to rostral and the brain sections containing the DVC and Midbrain were collected and kept in a 24 well plate containing phosphate buffered saline. These sections were then stored in a cryoprotectant solution (30% v/v glycerol, 30% v/v ethylene glycol in 1x phosphate buffered saline) at -20°C until they were ready to be stained using immunofluorescence. For staining, all sections were first washed 3x for 15 minutes in PBS at room temperature. All sections were then blocked and permeabilized using a 0.3% Triton-X, 10% BSA in PBS solution at 37°C for 1 hour. Following blocking and permeabilization, the DMV sections were stained for cholinergic neurons (anti-ChAT), phosphorylated alpha synuclein (anti-pSer129), and microglia (anti-IBA1, see Table for details) diluted in 0.1% Triton-X, 5% BSA in PBS overnight at 4°C on a rocker. The following day, sections were washed 3x for 20 minutes each at room temperature in PBS. The sections were then incubated with a donkey anti-goat AlexaFluor 488 antibody diluted in 0.1% Triton-X 3% normal donkey serum in PBS covered for 1 hour at room temperature on a rocker. The sections were then washed again in PBS and incubated with goat anti-guinea pig AlexaFluor 647 and goat anti-mouse AlexaFluor 568 secondary antibodies diluted in 0.1% Triton-X and 3% normal goat serum in PBS covered for 1 hour at room temperature on a rocker. Sections were then washed 3x in PBS again and counter stained with DAPI (1:4000), mounted, and coverslipped with Fluoromount Gold. Mounted slides were sealed with nail polish and stored in the dark until confocal imaging.

**Confocal Imaging**

Mouse brain and gut tissue was imaged using a Nikon A1 laser scanning confocal microscope using NIS Elements software (V5.0 Nikon). Samples were illuminated using 405, 488, 560, and/or 640nm diode lasers (Obis) through either a x20 (0.75NA Plan Fluor) mixed immersion, x40 (1.3NA Plan Apo) oil-immersion, or x60 (1.4NA Plan Apo) oil-immersion objective with emission detection set to 595/50, and 700/75nm, respectively. To avoid bleed-through, all images were acquired using channel series mode.

For imaging of mouse duodenal whole mounts, the myenteric plexus was first visualized and inspected. Any tissues with incomplete staining of the myenteric plexus were discarded. For remaining samples, 4-6 z-stack images were randomly taken across the sample with the total width of the myenteric plexus captured for each image. For pSer129 neuronal counts, images were taken 1.0 microns apart, for all other images, images were taken at 0.5-micron increments. For mouse brain imaging the central canal was first identified as a landmark for the DMV region. Large, stitched images, or multipoint acquisition was used to capture the right and left DMV (ChAT+ neurons on either side of the central canal). Z-stacks of 20-30 microns were taken at 0.5-micron increments.

Image processing was either performed in Fiji/ImageJ (see below) or, for representative images, in Nikon NIS Elements software. Representative images were background subtracted using the rolling ball algorithm and then denoised using the Elements Advanced Denoising feature set to a power of 2 for each channel.

**scRNA-Sequencing**

*Sample Preparation*

Single-cell RNA sequencing was carried out using the 10x Single-cell 3' Dual Index kit (v3.1). For each library, 2 animals were pooled. Animals were terminally anesthetized and then perfused with ice-cold PBS with 2% fetal bovine serum (FBS). The duodenum from each animal was dissected and any remaining mesentery was carefully removed on ice. The duodenum was then flayed open, cleaned, and finely cut into small pieces using a fresh razor blade. The duodenal pieces were then added to 5ml of Accumax dissociating agent and allowed to digest with mild agitation for 5 minutes at 34°C-37°C. Following digestion, the mix was further mechanically dissociated using a wide-bore P1000 pipette tip with gentle trituration. Fresh PBS+FBS was then added to the mix and the tissue was spun down at 340g for 3 minutes. After the first spin, the sample was then filtered through a 40μm tube-top filter and rinsed thoroughly with ice cold PBS+FBS. The sample was then spun down again under the same settings and the supernatant was aspirated. The remaining sample was re-suspended and counted under a microscope using a hemocytometer and Trypam Blue.

Once counted, cells were allocated to either enrichment for myeloid cells using CD11b magnetic bead sorting, or immunolabeling for CD45 to capture all gut immune cells. Myeloid enrichment was accomplished using a CD11b magnetic bead MojoSort kit (Biolegend, see Table). 10 million cells were allocated to CD11b positive selection according to the manufacturer protocol. Briefly, cells were first incubated with the biotin-antibody cocktail for 15 minutes on ice. Cells were then washed by hand and spun down at 300g for 3 minutes. Next, labelled cells were allowed to incubate in 100μl of MojoSort Buffer with the streptavidin nanobeads on ice for 15 minutes. Cells were re-washed and spun again at 300g for 5 minutes and then resuspended in 1ml and placed in a magnetic separator column. 3 rounds of separation, salvage, and washing were performed to enrich for CD11b+ cells. Following magnetic enrichment, cells were then incubated in a 1:2000 DAPI mix to distinguish live/dead cells.

Simultaneously, 4-6 million cells were allocated for CD45 labelling. Briefly, cells were allowed to incubate with CD45-FITC at 0.5μl per million cells in FACS buffer for 30 minutes, covered and on ice. Cells were then washed with PBS and a 1:2000 DAPI mix was added to allow for Live/dead labelling. Both the CD11b-enriched, and CD45-labelled cell samples were then subjected to FACS using a BD FACSAria III. Importantly, cells were sorted on the slowest speed using a 100μm nozzle and sorted into pre-wetted microcentrifuge tubes filled with 100μl of PBS+FBS. For each sort, 50-80,000 cells were sorted. Following FACS, sorted cells were washed 3 times to discard any debris/dying cells and filtered once more using a P1000 filter tip pipette. Cells were then counted one final time and were ready to load onto the 10x chip.

*Library Construction and Sequencing*

Only samples with at least 80% final viability by Trypam Blue were selected for chip-loading and further library construction. For each library, the maximum number of cells was loaded into each well of the chip. GEM recovery and cDNA amplification was performed according to the 10x protocol. All libraries were QC'd using a bioanalyzer high-sensitivity chip before proceeding with library construction. A final QC was also conducted using a Bioanalyzer High Sensitivity chip prior to freeze-down or sequencing. Libraries were kept at -20°C until they were sent for sequencing using an Illumina NovaSeq 6000 at the University of Florida's Interdisciplinary Center for Biotechnology Research NextGen DNA Sequencing Core. All libraries were sequenced to a depth of 100,000 read pairs per cell, and all libraries were sequenced on a single flow cell to help minimize any potential batch effects.

**Fecal Output**

Fecal output assays were carried out in a flow hood. Mice were transferred from their home cage into individual, covered to-go cups for 30 minutes. At the end of the 30 minutes, the fecal pellets were collected and counted for each animal. Pellets were blotted dry to remove any urine and then weighed together.

**Image Analysis**

All image analysis was performed using Fiji/ImageJ software and associated plugins. Details for each type of analysis are described below.

*Neuronal pSer129 $\alpha$Syn Counts*

Images were first background subtracted using the rolling ball method available in ImageJ then denoised using the Pure Denoise plugin with default settings selected. The appropriate channels were then thresholded and masked using the Triangle filter in ImageJ with the Autodetect set for threshold selection. Composite images were then made my combining the masked Tuj1 channel with the masked pSer129 channel. Throughout the pre-processing steps, images were quality checked. Any errors at any point in the process resulted in the image or slice being discarded from analysis (whichever was appropriate). A maximum intensity projection image was created and the total number of ganglia was counted. Ganglia were defined as areas where multiple fiber tracts came together and there were visible soma at the junction. To identify pSer129+ soma, the z-stack was scrolled through manually and positive soma were visibly identified, marked, and counted. In order to count, soma must be positive within the z-area of the myenteric plexus (i.e., positive signal above or below was not counted) and approximately half or more of the soma has to be positive for pSer129. Additionally, the shape of the signal had to approximate a cell body (i.e., scattered punctae were not counted). The total number of pSer129+ neurons was then added up and divided by the total number of ganglia to yield the average number of pSer129+ soma per ganglia.

*Macrophage pSer129 punctae*

For macrophage pSer129 punctae analysis, the same pre-processing workflow was followed as for the neuronal counting analysis. Following background subtraction, denoising, and thresholding and masking, the Image Calculator function in ImageJ was used to select pixels that were positive for both MHCII and pSer129. A sum slices projection was taken of the resultant image and the Analyze Particles Function was used to count the number of punctae as well as the average size of punctae within the macrophages. The average size of punctae was the metric reported.

*Macrophage-Neuron Contact*

Following the same pre-processing steps, the image calculator was again used to identify the pixels that were positive for both Tuj1 and MHCII. Regions of interest were then drawn around each ganglia in the Tuj1 channel. The double positive area for MHCII and Tuj1 in each ROI was then recorded and taken as a percentage of the total Tuj1+ ganglia area. For PFF-injected mice, ganglia with pSer129+ neuronal soma were noted. The proportion contact area for each ganglia for every image was averaged to give the reported values per mouse.

*C1q Analysis*

Images were subject to similar pre-processing as described above with the exception that a more stringent rolling ball radius (10) was selected for the background subtraction. To analyze the C1q deposition on neurons, the total peripherin+ area for each image was recorded. The image calculator was used to identify pixels that were positive for both C1q and Peripherin. The total C1q+/Peripherin+ area in square pixels was then divided by the total Peripherin+ area and multiplied by 100 to yield the reported measure.

A similar process was followed to measure the average C1q+ area per macrophage. Following the same pre-processing steps, the image calculator was used to identify pixels positive for both

MHCII and C1q. Separately, a maximum intensity projection of the thresholded and masked MHCII image was used to count the total number of macrophages in the field of view. The count, macrophages had to have a defined morphology, be entirely within the field of view, and not overlap with one another. The total C1q+/MHCII+ area was divided by the total number of macrophages counted to give the reported metric.
To identify C1q+/pSer129+ punctae within macrophages, the image calculator was again used with two iterations. The first was used to identify pixels that were double positive for both C1q and pSer129, the second was used to identify pixels positive for these two markers and MHCII. A sum slices projection was then made and the analyze particles function in ImageJ was used to calculate the number of total particles and their average size. The total number of particles was divided by the total number of macrophages counted (as above) to yield the average number of punctae per macrophage.

*Sholl Analysis*
Z-stack images were acquired using 60x oil immersion objective on a confocal laser microscope equipped with a Nikon A1 system. The DMV region was defined by the presence of positive ChAT neurons. Morphometric analysis was performed using Image J software. Maximum intensity projections were thresholded to create a binary image. Individual microglia were then isolated in the image and used to perform sholl analysis. For Sholl analysis, a line was drawn from the center of the soma to the end of the longest branch. Deprecated sholl analysis on the Neuroanatomy plug-in then drew concentric circles from the start to the end of the line at 1 μm increments and counted the intersections at each radii.

*Co-localization analysis*
Co-localization analysis was performed using Nikon Elements NIS analysis software. Images were pre-processed by rolling ball background subtraction and advanced denoising. The co-localization analysis option was performed through each slice of the z-stack and the Pearson’s R for each slice was averaged to give a co-localization value for the image.

**scRNA-Sequencing Analysis**
Following sequencing, fastq files were passed through the CellRanger (v6.1.1) pipeline for alignment, mapping, barcode assignment and UMI counting. The default mouse genome in the CellRanger pipeline was used to map reads. Libraries were visually inspected individually for quality control. Any library where <70% of reads could be confidently mapped to cells was not used for downstream analysis. Libraries from the same condition were then aggregated using the CellRanger aggregation function available on their cloud platform. The resulting barcode and count matrices were then passed through to Seurat v4.0[6] for downstream analysis.
Files were read into Seurat and underwent further quality control based off of the feature and UMI counts, and the mitochondrial contamination. The minimum and maximum feature counts were set to 200 and 6500, respectively while the percent mitochondrial-derived RNA was capped at 15%. Only cells that met these requirements proceeded to the next step of analysis. The data were then normalized, and the top 2500 highly variable features were identified. The saline and PFF libraries were then integrated using the ‘FindIntegrationAnchors’ and ‘IntegrateData’ functions in Seurat. The resultant combined dataset was then used for the remaining analyses. For an initial clustering analysis, the data were first scaled, then PCA (npcs=10) followed by dimensionality reduction via uniform manifold approximation and projection (10 dimensions) were performed. The FindNeighbors function (10 dimensions) and the FindClusters function (with resolution of 0.1) were then carried out. The default settings for all of the above functions were used except where specified. For DEG identification, the FindMarkers function was used to compare the gene counts in the Mp clusters or the T-cell clusters from the PFF vs the Saline cells.

To perform a high-resolution clustering analysis specifically on the macrophages, the "Mps" cluster was extracted from the larger dataset. These data were then renormalized and a new set of 2500 highly variable features were found. The Mp dataset was then re-scaled and a new PCA was performed (10 components). The number of components was determined by visual inspection of an elbow plot. Finally the data were reclustered with a higher resolution of 0.7. To analyze proportions, the total number of cells from each category in each cluster were tabulated and then either a z-test or Fisher exact test were performed to confirm statistical significance. For comparison of signatures from clusters of interest with published datasets, all significant (adjusted p-value <0.05) marker genes for that cluster were pulled into a list and compared against the published signature for disease associated microglia (DAM) or disease inflammatory macrophages (DIM) from the supplemental table of Silvin et al 2022. For functional annotation, the significant marker genes were entered into the Gene Ontology Panther Database and the resultant terms were ranked on the Fold Enrichment. To identify DEGs between Mp6 vs Mp2, the FindMarkers function was used to directly compare the two clusters. Again, the significant DEGs following p-value adjustment were input into the Gene Ontology Panther Database for functional annotation. A similar process was carried out for the T-cell re-clustering and subsequent analysis.

*Trajectory analysis*

Trajectory analysis was performed using the Monocle (v3) package in R[7]. Macrophage clusters identified via Seurat were loaded and re-clustered using the cluster_cells() functions with a resolution of 0.01. The resulting UMAP identified Mp2 and Mp6 as belonging to the same partition. This partition was then selected and the learn_graph() function followed by the order_cells() and root_cells() functions were used to develop a trajectory through pseudotime for the partition.

**Statistics**

Statistical analysis for scRNA-seq data was all performed in R Studio using the default settings of the functions described above. All other statistical analysis was performed in Graphpad Prism v9.0. Details for individual statistical tests can be found in the legend of each figure. For all datasets, normality was checked by visual inspection of a Q-Q plot. Bar graphs show the group mean +/- SEM unless otherwise stated.

**Supplemental Figure 1: Supporting data for Figure 1. Confocal images of the myenteric plexus.** (A) Thresholded and masked confocal images of the myenteric plexus in 30dpi mice. (B) Quantification of area covered by myenteric neurons at 30dpi expressed as a percentage of saline control showing no significant difference in myenteric plexus area between groups. Unpaired t-test, n=5 animals/group. (C) Thresholded and masked confocal images of myenteric macrophages at 30dpi. (D) Quantification of the area covered by myenteric macrophages at 30dpi expressed as a percentage of saline control showing no significant difference between groups. Unpaired t-test, p=0.9372, n=5 animals/group. (E) 20x images of the DMV (top), SNc (middle), and VTA (bottom) in PFF-injected mice at 30dpi stained for pSer129 α-synuclein showing no pathology in these nuclei at this time point. Images representative of experiments repeated in 5 animals and two independent experiments. Inset shows 4x image of region. (F) Sholl analysis of microglia in the DMV at 30dpi in non-injected, saline-injected, and PFF-injected mice (left) and corresponding area under the curve quantification (right) showing that gut PFF injection significantly increased the morphologic complexity of DMV microglia. One-way ANOVA (p<0.0001) with Dunnett's test for multiple comparisons. (G) Representative confocal images of the myenteric plexus in non-depleted and depleted mice at 7 days post-depletion showing effective macrophage depletion at 7 days post-depletion point. (H) Images of cortical microglia in non-depleted and depleted mice at 7 days post depletion showing preservation of microglia with this depletion strategy. (I) Thresholded, masked, and extracted myenteric macrophages from 30dpi saline and 30dpi PFF mice used for morphology analysis. Red outline indicates projection area. Quantification of the projection area (J), the MHCII intensity (K), and the area: perimeter ratio showed no significant difference between PFF-injected mice and saline controls at 30dpi. Mann-Whitney test, p=0.6905 (J), p=0.8413 (K), p=0.6905 (L), n=5 animals/group.

**Supplemental Figure 2: Supporting data for Figure 2. Experimental Design and Flow cytometry gating strategy for single-cell RNA sequencing.** (A) Experimental design and gating strategy for isolation of intestinal immune cells for droplet-based single-cell RNA sequencing. (B) U-MAP of 27,648 high-quality single-cell transcripts from mice 30 days post-injection with either saline or PFFs clustered into 13 transcriptionally distinct cell types. (C) Dot plot of cell clusters in (B) showing the top 2 unique marker genes for each cluster. (D) Feature plots indicating enrichment of marker genes *Cx3cr1, Lyz2,* and *Aif1* in the macrophage cluster. (E) U-MAP of T-cells extracted from the UMAP in (A) revealing 11 transcriptionally distinct sub-clusters. (F) Volcano plot of differentially expressed genes in saline vs. PFF T-cells. (G) Composition of the T-cell subclusters.

**Supplemental Figure 3: Enteric neuronal α-syn localizes to pre-synaptic regions.** (A) Representative confocal images of myenteric plexus (peripherin, pink) showing co-localization with α-syn (stained with an N-terminal monoclonal antibody clone 3H19, cyan) and pre-synaptic marker bassoon (yellow). (B) Quantification of co-localization between α-syn and nuclei (DAPI), general neuronal structures (peripherin), or pre-synaptic spaces (bassoon) using Pearson's co-localization coefficient showing that α-syn is highly co-localized with neuronal structures and has a higher co-localization with pre-synaptic markers specifically (one-way ANOVA $p<0.0001$ with Tukey's test for multiple comparisons). (C) Representative confocal images of α-syn (cyan) marked with an independent polyclonal antibody colocalizing with bassoon (yellow). (D) Quantification of co-localization between α-syn and nuclei (DAPI) vs α-syn and pre-synaptic spaces (bassoon) again showing preferential localization to the pre-synaptic areas (unpaired t-test $p<0.0001$). (E) Representative confocal images of myenteric plexus (peripherin, pink) showing co-localization between α-syn (stained with an N-terminal monoclonal antibody clone 3H19, cyan) and post-synaptic marker PSD95 (yellow). (F) Quantification of co-localization of α-syn with either peripherin or PSD95 showing no significant difference (unpaired t-test).

#### Supplemental Figure 1

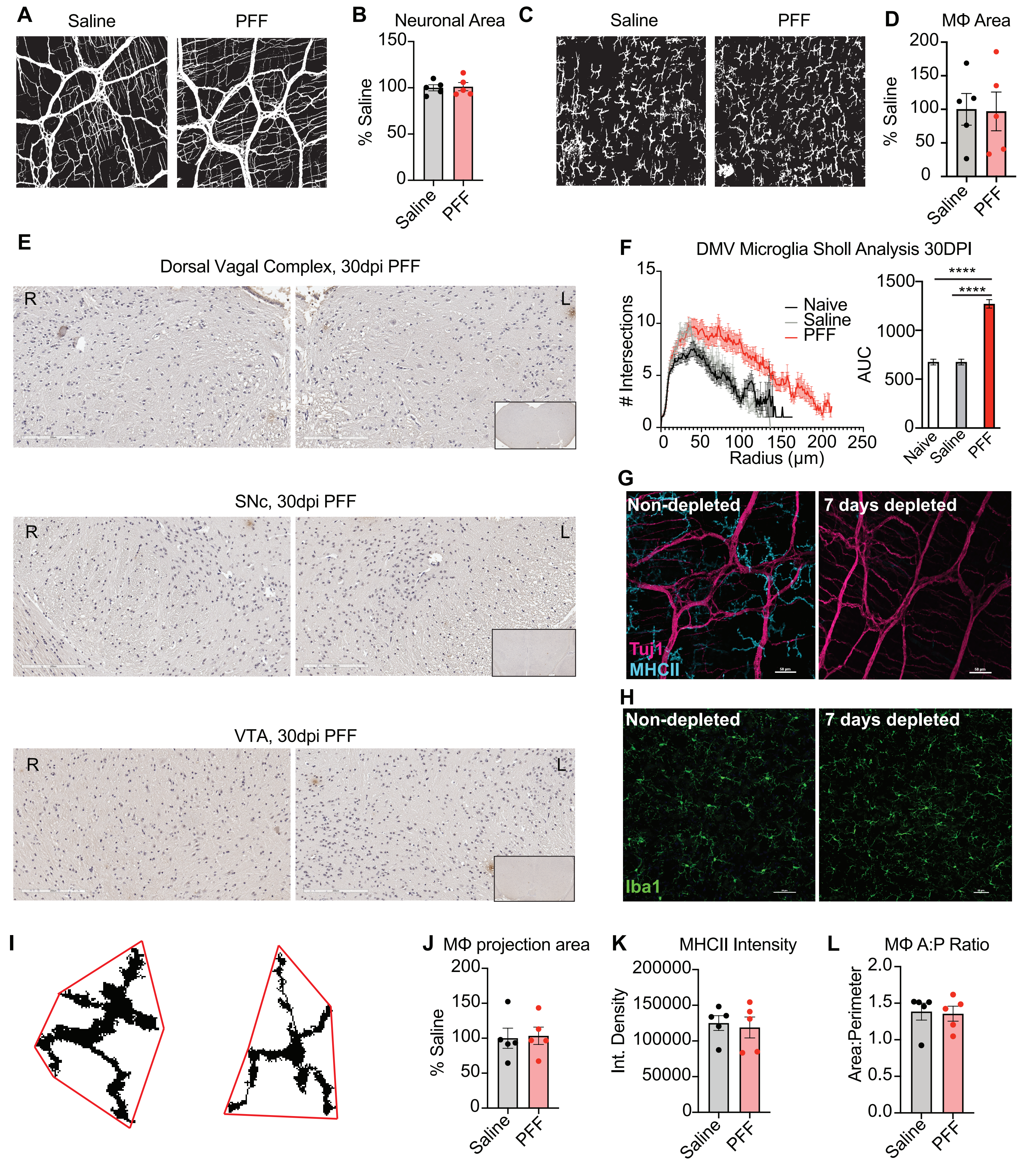

### Supplemental Figure 2

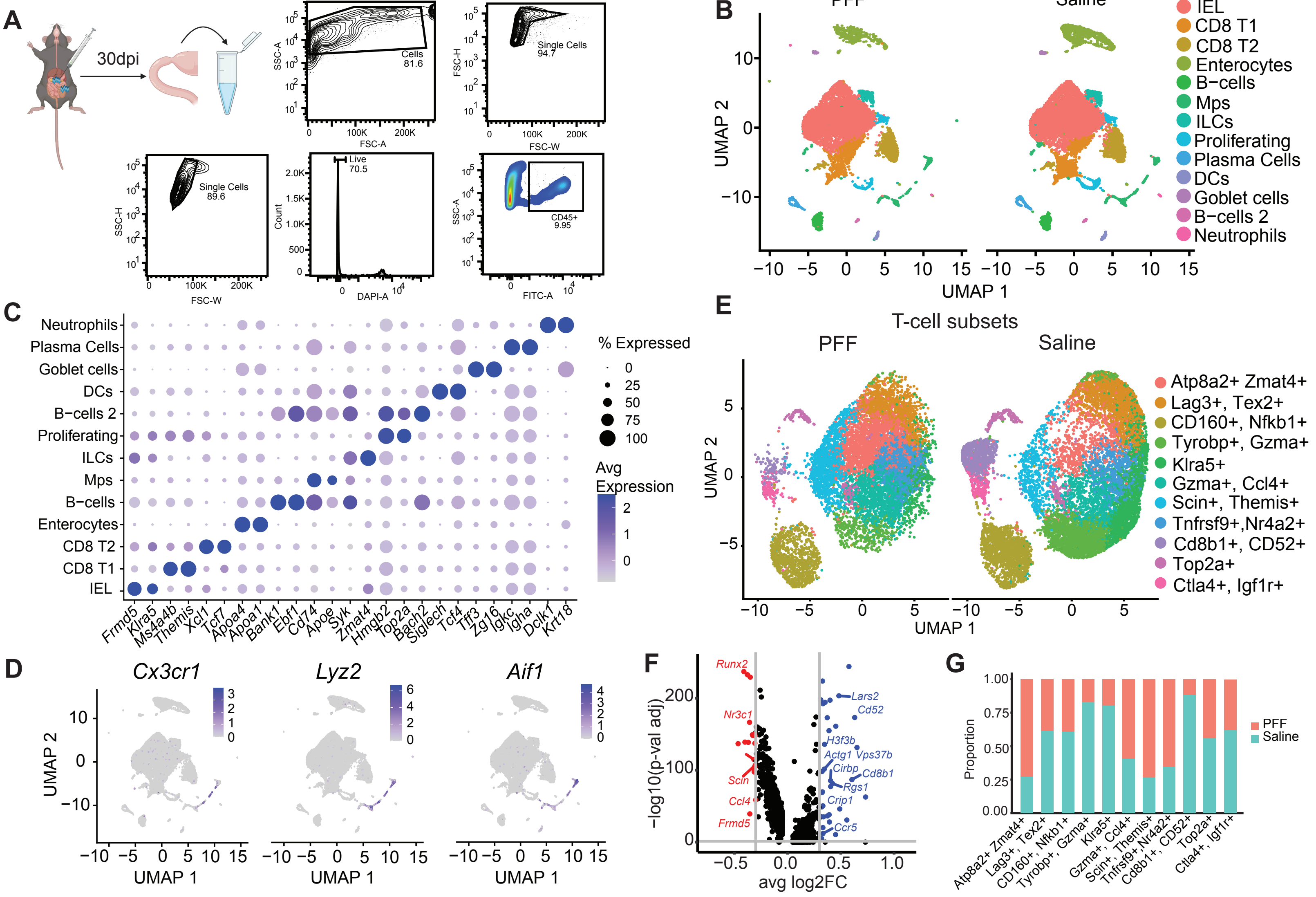

#### Supplemental Figure 3

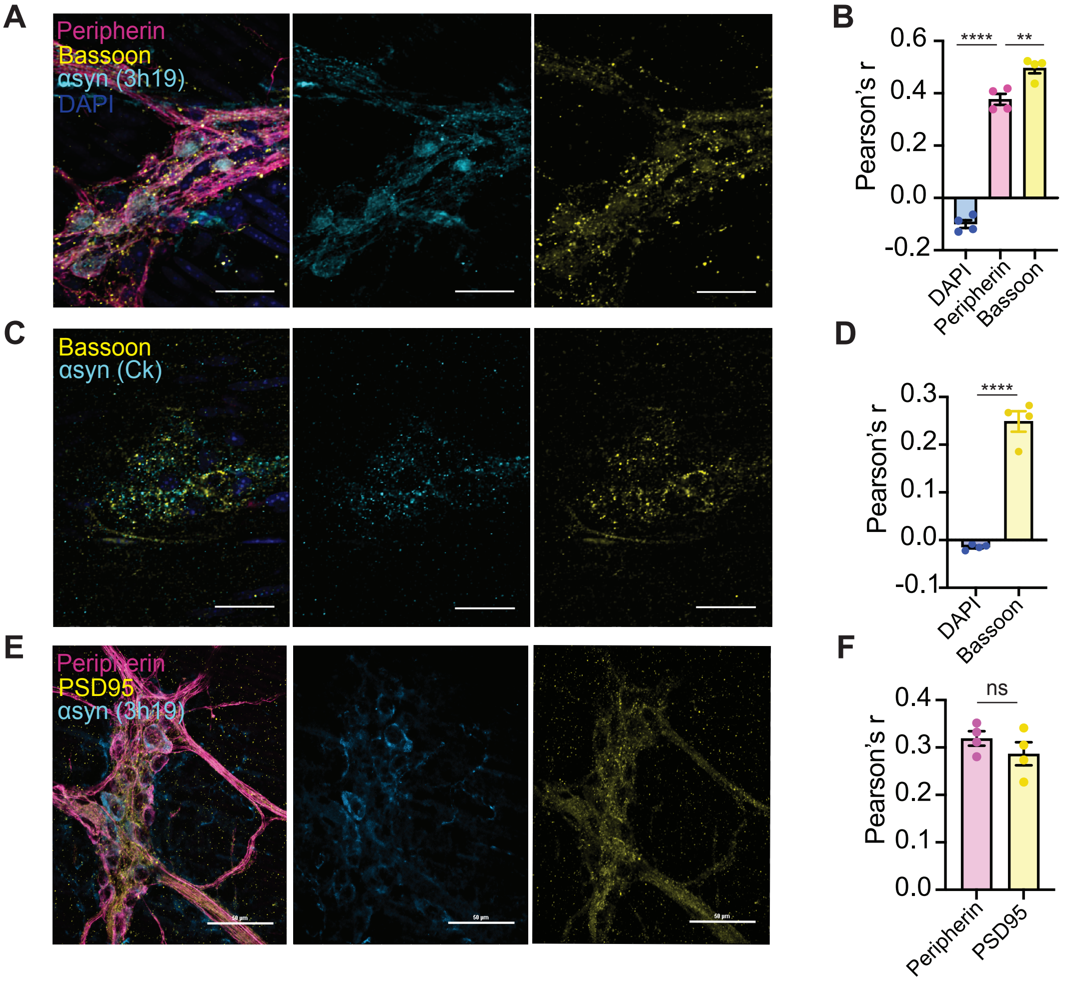
